# “UNTARGETING” AUTOANTIBODIES USING GENOME EDITING, A PROOF-OF-CONCEPT STUDY

**DOI:** 10.1101/2022.10.29.514381

**Authors:** Gerson Dierley Keppeke, Larissa Diogenes, Kethellen Gomes, Luis Eduardo Coelho Andrade

## Abstract

Autoantibodies are useful biomarkers of autoimmune diseases and some have direct pathogenic role. Current standard therapies for elimination of specific B/plasma-cell clones are not fully efficient. In this proof-of-concept study, we used the CRISPR/Cas9 genome-editing system to knockout V(D)J rearrangements that produce pathogenic autoantibodies *in vitro*.

HEK293T cell lines were established with stable expression of two monoclonal antibodies, a humanized anti-dsDNA (clone 3H9) and a human-derived anti-nAChR-α1-subunit (clone B12L). For each clone, five CRISPR/Cas9 guided-RNAs (T-gRNAs) were designed to target the heavy chain CDR2/3 variable regions. After CRISPR/Cas9 editing, levels of secreted immunoglobulins were evaluated, in addition to 3H9 anti-dsDNA reactivity by ELISA and B12L anti-AChR reactivity using cells overexpressing mouse genes of AChR-α1/β1/δ/γ/ε-subunits.

The T-gRNAs decreased the expression of the heavy chain to ∼50-60%, compared to >90% in Non-Target-gRNA. Levels of secreted IgG and reactivity to the respective target antigens decreased ∼90% and ∼95% after knockout with the T-gRNAs compared to Non-Target-gRNA for clones 3H9 and B12L, respectively. Sequencing indicated the presence of *indels* at the Cas9 cut-site, which could lead to codon jam, the likely cause of the knockout. Additionally, remaining secreted 3H9 antibodies presented variable reactivity to dsDNA among the five T-gRNA, suggesting that the exact Cas9 cut-site and *indels* may further interfere with antibody-antigen interaction.

CRISPR/Cas9 genome-editing was very effective to knockout the Heavy-Chain-IgG genes, considerably affecting the secretion and binding capacity of the autoantibodies *in vitro*, warranting application of this concept to *in vivo* models as a potential novel therapeutic approach for autoantibody-mediated diseases.

**Highlights:**

➢ Autoantibodies can have a direct pathogenic role in some autoimmune diseases.
➢ Elimination of specific B/plasma-cell clones is not attainable with current therapies.
➢ CRISPR/Cas9 allows targeting of specific DNA sites, such as V(D)J rearrangements.
➢ CRISPR/Cas9 genome-editing was very effective in knocking out the heavy chain of autoantibodies.
➢ Indels introduced at Cas9 cut site interfered with autoantibody-antigen interaction.

## 1. Introduction

Autoantibodies are invaluable biomarkers of autoimmune diseases and have a direct pathophysiological role in several of them. Examples of pathogenic autoantibodies include anti-platelet autoantibodies in autoimmune idiopathic thrombocytopenia; antibodies against desmoglein-1 and 3 in pemphigus; thyroid-stimulating autoantibodies in Graves’ disease; and the antibodies against the nicotinic acetylcholine receptor (anti-nAChR) in Myasthenia Gravis (MG), that interfere with neural-muscular signal transmission [1]. A comprehensive review of the topic can be found elsewhere [1].

The antigenic specificity of an antibody is assembled during the development of B lymphocytes by a DNA recombination process called V(D)J joining. When the rearrangement that produces the variable regions of the light and heavy chains of an antibody is terminated and the affinity maturation completed, that will be the only type of antibody that will be produced by that B lymphocyte clone, now-differentiated into plasma cells, a B cell specialized in and devoted to producing antibodies [2]. Different antibodies target the same antigen, a phenomenon called polyclonality, and the repertoire of possible recombination configurations for a given antigen is limited at the individual level, although huge among individuals within the species [3, 4]. Autoantibodies are mainly polyclonal and, among the various clones for a given autoantigen, there is considerable heterogeneity in the avidity and affinity to the target antigen, with corresponding heterogeneity in their pathogenic potential. Of interest to the pathogenic potential of the autoantibodies, some clones “prefer” strategic areas of the autoantigen, such as the main immunogenic region (MIR) in the nAChR, a small portion of the extracellular part of subunit alpha 1, where the neurotransmitter acetylcholine binds to [5].

The major sources of high-affinity immunoglobulin type G (IgG) in adults are the long-lived plasma cells (LLPC). These cells do not undergo somatic hypermutation, therefore each clone can only produce a single class of immunoglobulin with an unchangeable antigenic specificity. Different from B-cells, LLPC express an insignificant amount of surface antibodies and do not depend on interaction with antigens for continuous secretion of large amounts of immunoglobulin. LLPCs are also considered truly long-lived, as their population does not depend on constant replenishment by memory B cell re-stimulation [6-8]. Current therapies focused on the depletion of B cells, such as anti-CD20 monoclonal antibodies, do not affect LLPC and therefore have little effect on the concentration of circulating high-affinity autoantibodies. Other types of immunomodulatory drugs, such as immunobiological drugs targeting other receptors, cytokines, or interfering with checkpoints, have also shown little effect on LLPC in clinical trials [9]. Currently, the most effective way of removing high amounts of circulating autoantibodies employs plasmapheresis or bone marrow transplantation. Plasmapheresis does not solve the problem, though, since autoantibody concentration returns to previous levels in a few weeks. Bone marrow transplantation has several potential problems affecting over 50% of patients, including a high risk of infection, chronic Graft versus Host Disease, and considerable morbi-mortality [10].

Ablation of specific B/plasma-cell clones in autoimmune diseases caused by pathogenic autoantibodies would greatly improve treatment, thus scientists worldwide have proposed a variety of strategies. We will mention two recent experimental strategies to achieve such an endeavor, although there are several others [11].

1. Chimeric autoantibody receptor T cell (CAART) was recently proposed as a way to deplete antigen-specific B cells. The antigen desmoglein 3 (DSG3), a target of autoantibodies in Pemphigus Vulgaris (PV), was fused to CD137-CD3ζ signaling domains. The construct was transduced into primary human T cells to generate the DSG3-CAART. These cells were injected into an adoptive mouse model of PV, being able to successfully eliminate B-cells producing anti-DSG3 and considerably decreasing the amount of circulating anti-DSG3 autoantibodies [12].
2. Another proposal makes the use of a technology called affinity matrix (AM), where the recombinant antigen of interest is chemically fused to an antibody against plasma cells surface markers CD138 or CD44. Once binding of the CD138/CD44 molecules at the surface of plasma cells producing antibodies to that antigen, the complex (anti-CD138+autoantigen) will be preferably targeted by the autoantibodies being produced by that cell. The Fc region of the autoantibody at the cell surface will recruit complement molecules resulting in a complement-dependent cytotoxicity “suicide”. To test the technology, AChR-CD138 AM was injected into mice previously double immunized with *Torpedo*-AChR, an active model of Myasthenia Gravis. The AM was able to significantly deplete AChR-specific plasma cells in vivo [13, 14].

Over the past 10 years, controlled gene-editing gained a variety of new possibilities with the advent of a new system, which uses nucleases such as Cas9 and Cas13 (among others) with the ability to cleave the double DNA strand in a specific region determined by a guiding RNA (gRNA). This system is called CRISPR/Cas9 (Clustered Regularly Interspaced Short Palindromic Repeats) [15]. Although many applications are possible with CRISPR/Cas9 [16], one of the most efficient is to knockout a specific gene. After cleavage by Cas9 of a specific gRNA-determined target site, repair enzymes try to reconnect the DNA double-strand, but random errors, such as nucleotide deletions or additions (*indels*), often occur, otherwise the gRNA/Cas9 will return and cut again until the sequence have been sufficiently altered by the *indels* to avoid the gRNA annealing. When the *indels* are multiple of three, amino acids can be removed or inserted altering protein function, however in most cells the *indels* are not multiple of three, codons are jammed and the translation of that gene is damaged permanently due to early stop codons, generating a knockout [16].

Elimination of specific B/plasma-cell clones is not attainable with current standard therapies. In this proof-of-concept study, our objective is to use the CRISPR/Cas9 genome-editing system to knockout V(D)J rearrangements that produce pathogenic autoantibodies *in vitro*. Adaptation of this method to *in vivo* modulation of pathogenic B/plasma-cell clones could pave the way to a novel targeted therapy platform in frame with the concept of precision medicine paradigm.

## 2. Results

### 2.1. Construction of antibody-producing cell-lines

Cell lines were established with stable expression of two human recombinant antibodies: 1) A human-derived anti-nAChR-α1-subunit, clone B12L; 2) A humanized anti-dsDNA, clone 3H9. The anti-AChR clone B12L was recently obtained by Makino *et*.*al*. [17] from a human patient with MG and demonstrated to induce pathogenic MG-like features in an *in vivo* passive transfer model of rat [17]. The complete variable region sequences for the heavy and light lambda chains were kindly shared by personal communication. The anti-dsDNA clone 3H9 was obtained over three decades ago (GenBank: M18234.1; M18237.1), derived from an MRL/lpr mouse, a murine model of SLE. It was demonstrated to bind both ssDNA and dsDNA [18].

The variable regions of the heavy chain (VH) genes of B12L and 3H9 antibodies were cloned into a vector (plasmid 1) that contains the human immunoglobulin heavy chain constant region, plus a reporter fluorescent gene (iRFP670) interleaved by T2A (**pCMV_VH-*h*IgG1-Fc_T2A_iRFP670)** (Figure 1 and Supp. Figure S1A). The variable regions of the light chain (VL) genes of both antibodies were cloned into vectors (plasmid 2) containing the Lambda (plasmid 2a) or Kappa (plasmid 2b) constant sequence plus the reporter eGFP, interleaved by T2A. Recombinant B12L uses lambda chain (**pCMV_VL-*Lambda-*CL_T2A-eGFP)** and recombinant 3H9 uses kappa chain **(pCMV_VL-*kappa-* CL_T2A-eGFP)** (Figure 1 and Supp. Figure S1A). After simultaneous transfection of both vectors for each recombinant antibody (VH + VL-L/k), the HEK293T cells were selected and sorted (Supp. Figure S1B). Over 90% of the cells showed stable expression of both chains (Figure 1B-C). These two antibody-expressing cell lines were used throughout the study.

**Figure 1.**
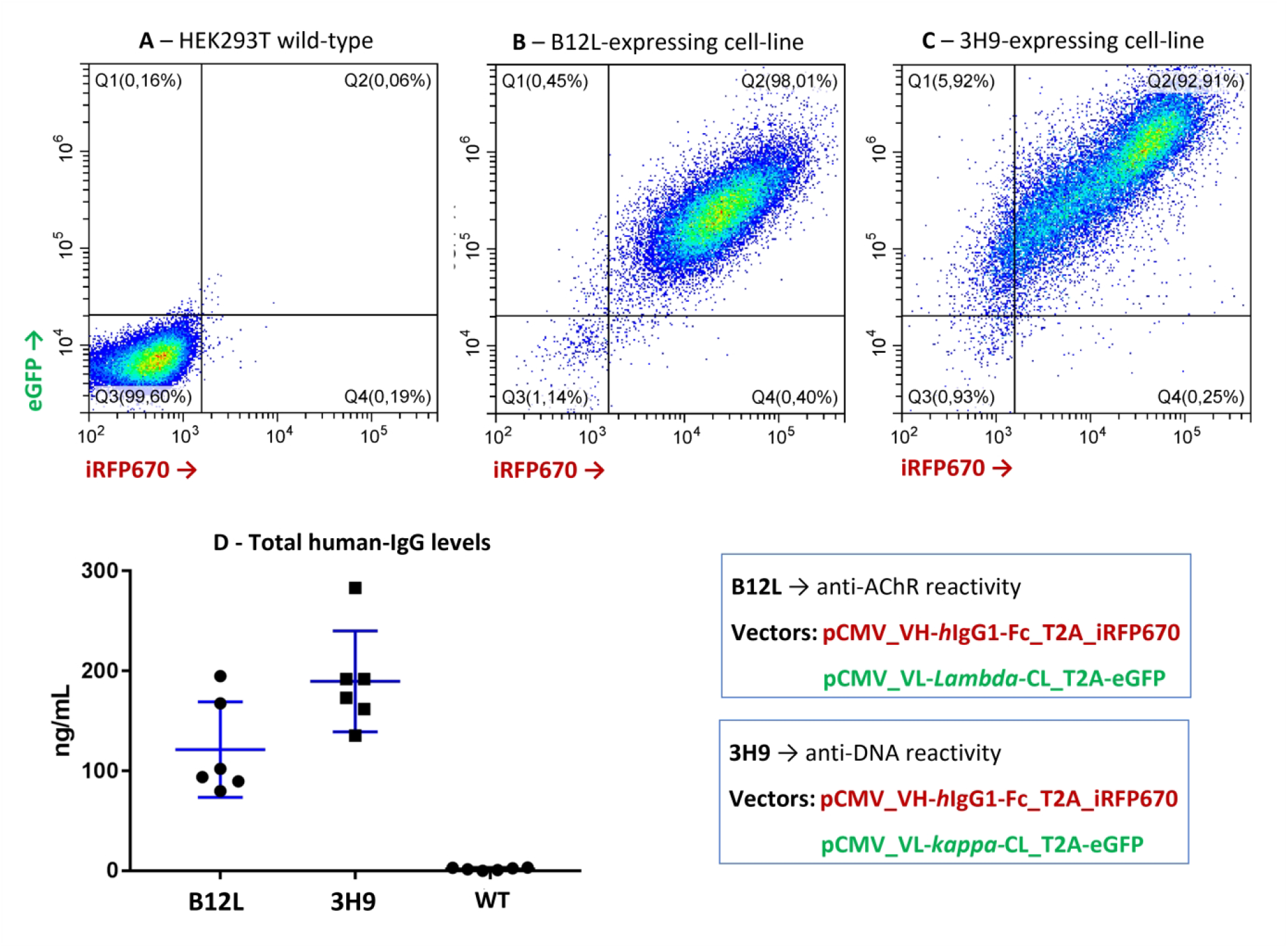
Phenotypic analysis of the recombinant antibody-expressing cell lines. Two vectors carrying the heavy (VH-Fc) or the light (VL) chains were constructed for each antibody. Each plasmid contains a fluorescent reporter molecule (eGFP or iRFP670) interleaved by a 2A sequence. **(A)** Wild-type HEK293T. **(B-C)** Cells were transfected with both plasmids, after selection and sorting, over 90% of the cells showed stable expression of both chains (Q2 = quartile 2 in B and C). In graphs (A-C), 30k events are shown. **(D)** Total recombinant human IgG levels were measured in the culture supernatant by ELISA. Supernatant was collected from the B12L-expressing cells and 3H9-expressing cells or wild-type (WT) (n=6 collections for each cell construct). Error bars = S.D.

### 2.2. Reactivity of the secreted recombinant antibodies

To quantify the recombinant antibody production, total human IgG was determined in culture supernatant (n=6) from the antibody-expressing and control cells, respectively (see methods for details). B12L-expressing cells secreted ∼120 ng/mL of immunoglobulins on average. 3H9-expressing cell-line secreted ∼190 ng/mL on average. (Figure 1D).

To test the reactivity of the recombinant antibodies to its respective targets, specific assays were applied. The reactivity of B12L against the acetylcholine receptor (AChR) was tested using HEK293T cells were transfected with two plasmids encoding the five subunits of mouse nicotinic AChR (α, β, δ, γ and ε) plus Rapsyn (HEK293T-mAChR) (Figure 2A). These cells were used as substrate in a cell-based indirect immunolabeling reaction where the B12L-containing supernatant was used as the primary probe and an anti-human IgG conjugated to FITC was used as the secondary probe. HEK293T-mAChR immunolabeled by B12L was analyzed by cytometry or microscopy (Figure 2A-D and Supp. Figure S2). Since the construct also contains rapsyn, the AChR subunits assemble into clusters at the cells’ surface [19] and these clusters are labeled by the recombinant B12L (arrows in Figure 2D). An average of ∼80% of HEK293T-mAChR were labeled by B12L, compared to <20% of wild-type HEK293T cells. Supernatant collected from cells that do not secret any antibody (also wild-type) labeled <20% of HEK293T-mAChR cells. Therefore, the secreted recombinant B12L specifically binds AChR in the cell-based assay. The reactivity of 3H9 against dsDNA was evaluated by a quantitative ELISA, yielding an average of ∼850 international units (IU/mL) obtained in the supernatant of six cultures (Figure 2F). In the HEp-2 IFA analysis, the 3H9 recombinant antibody presented a homogeneous nuclear pattern, similar to human serum with anti-dsDNA autoantibody. The average titer of 3H9 supernatant in the HEp-2 IFA was 1/40. (Figure 2G-J). These results confirm 3H9 reactivity to dsDNA.

**Figure 2.**
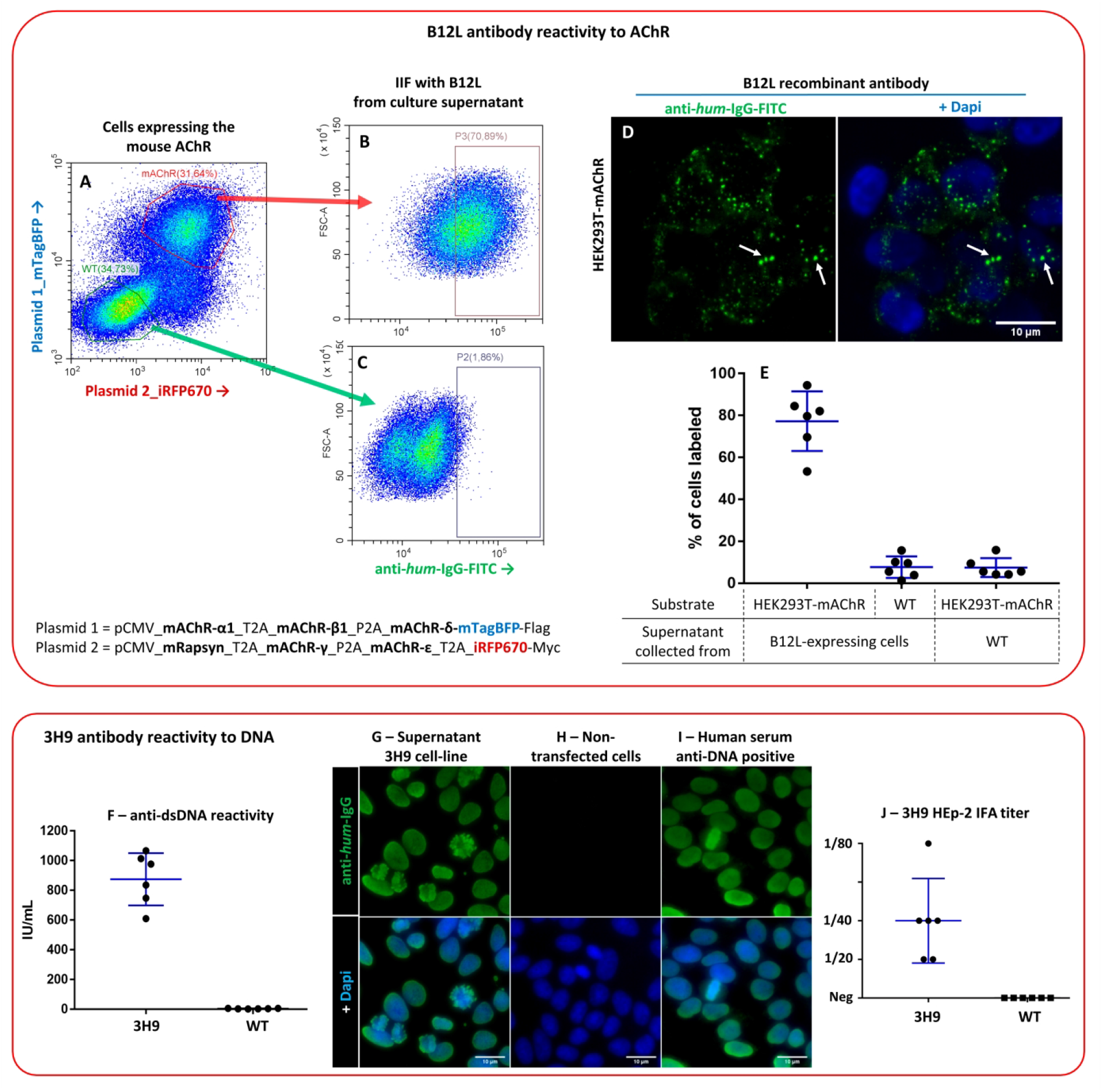
Anti-AChR reactivity by B12L and anti-DNA reactivity by 3H9 antibodies. **(A-E)** HEK293T cells were transfected with two plasmids encoding the five subunits of mouse nicotinic AChR plus Rapsyn (HEK293T-mAChR cells). **(A)** Flow cytometry analysis shows that ∼30% of the cells express the reporter genes for plasmid 1 (mTagBFP) and plasmid 2 (iRFP670). **(B-D)** HEK293T-mAChR cells **(B and D)** and WT HEK293T cells **(C)** were used as substrate for indirect immunofluorescence (IIF) with supernatant from B12L antibody-producing cells as primary probe. Cells were analyzed by flow cytometry **(B-C)** or by microscopy **(D)**. Arrows in **(D)** indicate AChR clusters, labeled by recombinant B12L plus anti-human IgG conjugated to FITC (green). **(E)** Proportion of HEK293T-mAChR or wild-type cells labeled by B12L supernatant, analyzed by flow cytometry, for 6 supernatant batches. **(F, G, H, J)** Supernatant from 3H9 antibody-producing cells (n=6) or wild-type (WT) cells were tested for anti-dsDNA reactivity by ELISA **(F)** or by HEp-2 immunofluorescence assay (HEp-2 IFA) **(G-I). (G)** The 3H9 recombinant antibody presents a homogeneous nuclear pattern in the HEp-2 IFA, **(I)** similar to human serum with anti-DNA autoantibody. **(J)** The average HEp-2 IFA titer was 1/40. Error bars = S.D.

### 2.3. CRISPR-Knockout of the VH of the recombinant antibody genes

Using the guided RNA designing tool CHOPCHOP v3 [20], five targeted gRNAs (T-gRNA) were designed for each antibody VH: two targeting the CDR2 and three targeting the CDR3 of the B12L-VH gene, and two targeting the CDR2 and three targeting the CDR3 of the 3H9-VH gene (Supp. Table S1). A non-targeted gRNA (NT-gRNA) was used as control. To minimize the off-target effect, and therefore preserve specificity, we selected T-gRNAs with at least 2 mismatches, considering the location of the PAM (protospacer adjacent motif) sequence. The chosen T-gRNA targets were within or neighboring the CDR2/3 of the antibody VHs.

T-gRNAs were cloned into a plasmid containing a U6 promoter and the reporter fluorescent gene mTagBFP. Each T-gRNA expression vector was transfected together with another plasmid containing sp*Cas9*. To ensure that the cells received the CRISPR/Cas9 system, 24 hours after transfection, cells were sorted for the presence of the reporter mTagBFP. After five days, the culture supernatant was collected for analysis of the effect of the editing on the levels and reactivity of the secreted antibodies.

We first analyzed the effect of CRISPR editing on VH expression by determining the expression of the iRFP670 reporter gene (Figure 3). For B12L-VH, the expression of the reporter gene decreased to ∼60% on average among the cells transfected with each of the five T-gRNAs individually compared to >95% of cells expressing iRFP670 in NT-gRNA-transfected cells. The lowest expression rate was observed for B12L-T-gRNA-4 (55%), which targets B12L-CDR3. Transfecting the two gRNAs targeting CDR2 together or the three gRNAs targeting CDR3 together produced a comparable decrease in VH gene expression, ∼60% and ∼55%, respectively (Figure 3C and D). Intracellular levels of IgG measured by ELISA also decreased to ∼55% on average for cells transfected with T-gRNA-4 and T-gRNA-5 targeting B12L-VH-CDR3, when compared to intracellular levels of IgG in cells transfected with NT-gRNA (Supp. Figure S3A). For the 3H9-VH gene, the expression of the reporter iRFP670 decreased to ∼53% on average among the cells transfected with each of the five T-gRNAs individually, compared to >94% in cells transfected with NT-gRNA. The lowest expression rate was observed for cells transfected with 3H9-T-gRNA-5 (45%) that targets 3H9-CDR3. Transfecting cells with the two gRNAs targeting CDR2 together or the three gRNAs targeting CDR3 together produced a comparable decrease in VH gene expression, ∼58% and ∼47%, respectively (Figure 3J and K). Intracellular levels of IgG decreased even further to ∼25% on average for cells transfected with T-gRNA-4 and T-gRNA-5, both targeting 3H9-VH-CDR3, when compared to intracellular levels of IgG in cells transfected with NT-gRNA (Supp. Figure S3B).

**Figure 3.**
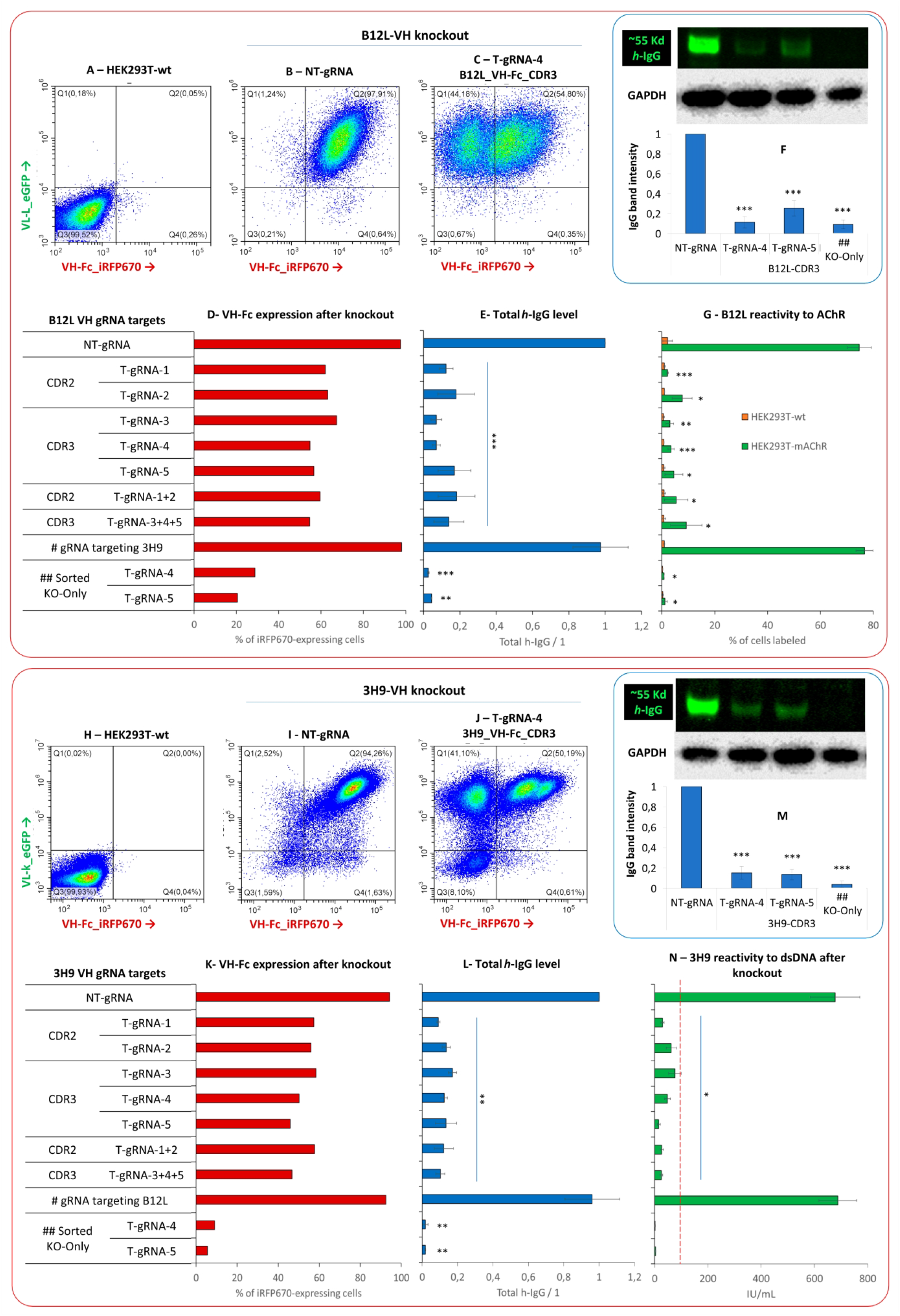
Knockout of VH with CRISPR/Cas9. Five T-gRNAs were designed for each antibody VH gene, two targeting the CDR2 and three targeting the CDR3 of the B12L-VH gene, and two targeting the CDR2 and three targeting the CDR3 of the 3H9-VH gene. A non-targeted gRNA (NT-gRNA) was used as control. **(A and H)** Wilde-type HEK293T. **(B-C)** B12L-expressing cells targeted by CRISPR. **(I-J)** 3H9-expressing cells targeted by CRISPR. **(B-D and I-K)** Five days after editing, cells were analyzed for the expression of the reporter gene (iRFP670) from B12L-VH-Fc or 3H9-VH-Fc, showing a decrease to ∼55% on average for all T-gRNAs in both antibody-producing cell lines. **(E-G and L-N)** Culture medium was collected on day 5 (n=4) and heavy chain IgG (*h*-IgG) was analyzed by ELISA (**E and L**) and by immunoblot (**F and M**); for both analyses, the concentration or band intensity obtained with NT-gRNA-transfected cells was defined as 1 in each batch of culture medium collected. Total cell extract was loaded in the gel for GAPDH labeling (**F and M**). **(G)** Reactivity of B12L against AChR determined by immunolabeling using HEK293T-mAChR as substrate and analysis by flow cytometry. **(N)** Reactivity of 3H9 against dsDNA determined by ELISA; the red dotted line at 100 IU/mL indicates the manufacturer’s cut-off recommendation for positive/negative determination in patient samples. **#** gRNA targeting 3H9-VH-Fc shows no effect on B12L-VH-Fc expression **(D-G)** and gRNA targeting B12L-VH-Fc shows no effect on 3H9-VH-Fc expression **(K-N)**. **##** Sorted “knockout-only (KO)” cells (check Supp. Figure S3). Error bars = S.D. in (E-M) and S.E.M. in (N). Statistics, p values for each T-gRNA compared to NT-gRNA by paired *t* Student test: *p<0.05; **p<0.01; ***P<0.001.

Altogether, this data indicates that the T-gRNAs found their intended targets with various efficiencies, since the frequency of cells expressing the VH reporter iRFP670 varied from 40% to 60%, and the levels of intracellular IgG varied from 25% to 50% with the different T-gRNAs. Of note, expression of the light chain (VL-Lambda/Kappa-CL_T2A-eGFP) was not affected in any of the T-gRNAs, as indicated by the GFP reporter (Figure 3C and J).

### 2.4. CRISPR-Knockout affects secretion and reactivity of the recombinant antibodies

In the collected supernatants of transfected cells, we first analyzed total human IgG by ELISA and western blot as indicative of the effect of the editing in the amount of antibody secreted by the cells. For both analyses, in each batch of culture medium collected (n=4), concentration found in NT-gRNA was normalized as 1 (Figure 3E-F and L-M). For B12L, IgG levels measured by ELISA decreased to ∼0.13 (∼87% of decrease) on average among cells transfected with the five T-gRNAs alone or grouped. The lowest levels occurred in cells transfected with B12L-T-gRNAs 3 or 4, targeting B12L-CDR3, both decreasing to 0.07 (total decrease of ∼93%) (Figure 3E). When analyzed by western blot, the *h*-IgG band intensity was ∼11% and ∼25% in cells transfected with T-gRNAs 4 and 5, respectively, compared to cells transfected with the NT-gRNA (Figure 3F). For 3H9, IgG levels decreased to ∼0.12 (∼88% of decrease) on average among cells transfected with the five T-gRNAs alone or grouped. The lowest level was observed for cells transfected with 3H9-T-gRNA-1 that decreased to 0.09 (∼91% decrease) (Figure 3L). By western blot, the *h*-IgG band intensity decreased to ∼15% and ∼13% in the supernatant of cells transfected with 3H9-T-gRNAs 4 and 5, respectively, compared to cells transfected with the NT-gRNA (Figure 3M).

Anti-AChR reactivity of B12L-secreted antibodies after CRISPR knockout was analyzed by immunolabeling assay with HEK293T-mAChR cells as substrate (Figure 2A-C). The proportion of HEK293T-mAChR labeled by B12L-knockout antibodies decreased to ∼5% on average for the cell lines transfected with each of the five T-gRNAs alone or grouped, compared to ∼75% for those transfected with the NT-gRNA (Figure 3G). The lowest proportion of HEK293T-mAChR cells labeled was observed for antibodies from cells transfected with B12L-T-gRNA 1 (targeting B12L-CDR2) and with B12L-T-gRNA 3 or 4 (both targeting B12L-CDR3) (∼3% for all three), which was similar to the proportion of labeling of HEK293T-wt by antibodies from cells transfected with B12L-NT-gRNA, thus these could be considered negative or undetectable (Figure 3G).

Anti-dsDNA reactivity for the secreted 3H9 after CRISPR knockout was analyzed by specific ELISA. Among cells transfected with the five 3H9-VH-T-gRNAs alone or grouped, the average anti-dsDNA reactivity was ∼40IU/mL, significantly below the ∼680IU/mL observed for cells transfected with the NT-gRNA. The lowest anti-dsDNA reactivity was observed for cells transfected with 3H9-T-gRNA-5, ∼15 IU/mL, representing a decrease of ∼98% when compared to cells transfected with NT-gRNA (Figure 3N).

Altogether, these data indicate that IgG secretion decreased over 90% after CRISPR knockout of B12L and 3H9 VH genes. Reactivity to the respective antigens was also significantly affected for both antibodies, down to virtually undetectable levels for some T-gRNAs, such as B12L-T-gRNAs 3 and 4, and 3H9-T-gRNA 5. As expected, gRNA targeting to 3H9-VH-Fc showed no effect on B12L-VH-Fc expression and gRNA targeting to B12L-VH-Fc showed no effect on 3H9-VH-Fc expression (Figure 3).

The effects of genome editing on the expression of VH-*h*IgG1-Fc could be monitored by the reporter gene iRFP670. Thus, five days after transfection of the CRISPR knockout system, cells not expressing iRFP670 were sorted, resulting in a population of cells with “highly effective” VH-Fc knockout (“KO-Only” cells) (Supp. Figure S4). KO-Only cell lines were obtained using B12L-T-gRNAs 4 and 5, and 3H9-T-gRNA 4 and 5. For B12L-KO-Only cells, total *h*-IgG measured by ELISA was down to ∼0.03 (97% decrease) and by western blot the band intensity was decreased >90% (*h*-IgG 55Kd band visually undetectable) compared to cells transfected with NT-gRNA (Figure 3E-F). B12L-KO-Only supernatant reactivity to AChR was also considered undetectable (Figure 3G). For the 3H9-KO-Only cells, total *h*-IgG measured by ELISA was down to ∼0.01 (99% decrease) and also visually undetectable by western blot, and the 3H9-KO-Only *h*-IgG 55Kd band intensity decreased 97% compared to cells transfected with NT-gRNA (Figure 3L-M). Reactivity to dsDNA by 3H9-KO-Only was also considered undetectable (Figure 3N).

### 2.5. VH s*equencing show indels at Cas9 cut-site*

The heavy chain variable (VH) regions containing the gRNAs targeting sites were amplified by PCR and sequenced forward and reverse using the Sanger method. For both antibodies, transfection with T-gRNAs deranged the sequence starting from the Cas9 cut-site, 3 bases from PAM, forward and reverse, suggesting insertion or deletions (*indels*) in that region. An example is provided in Supp. Figure S5A, where the sequence in the Cas9 cut-site of B12L-VH-CDR3 is deranged in cells transfected with B12L-T-gRNA-4. The same was observed for the 3H9-knockout, with derangement of the sequence forward and reverse at Cas9 cut-site after edition of CDR3 region with 3H9-T-gRNA-4 or 3H9-T-gRNA-5 (Figure 4B and Supp. Figure S5B).

**Figure 4.**
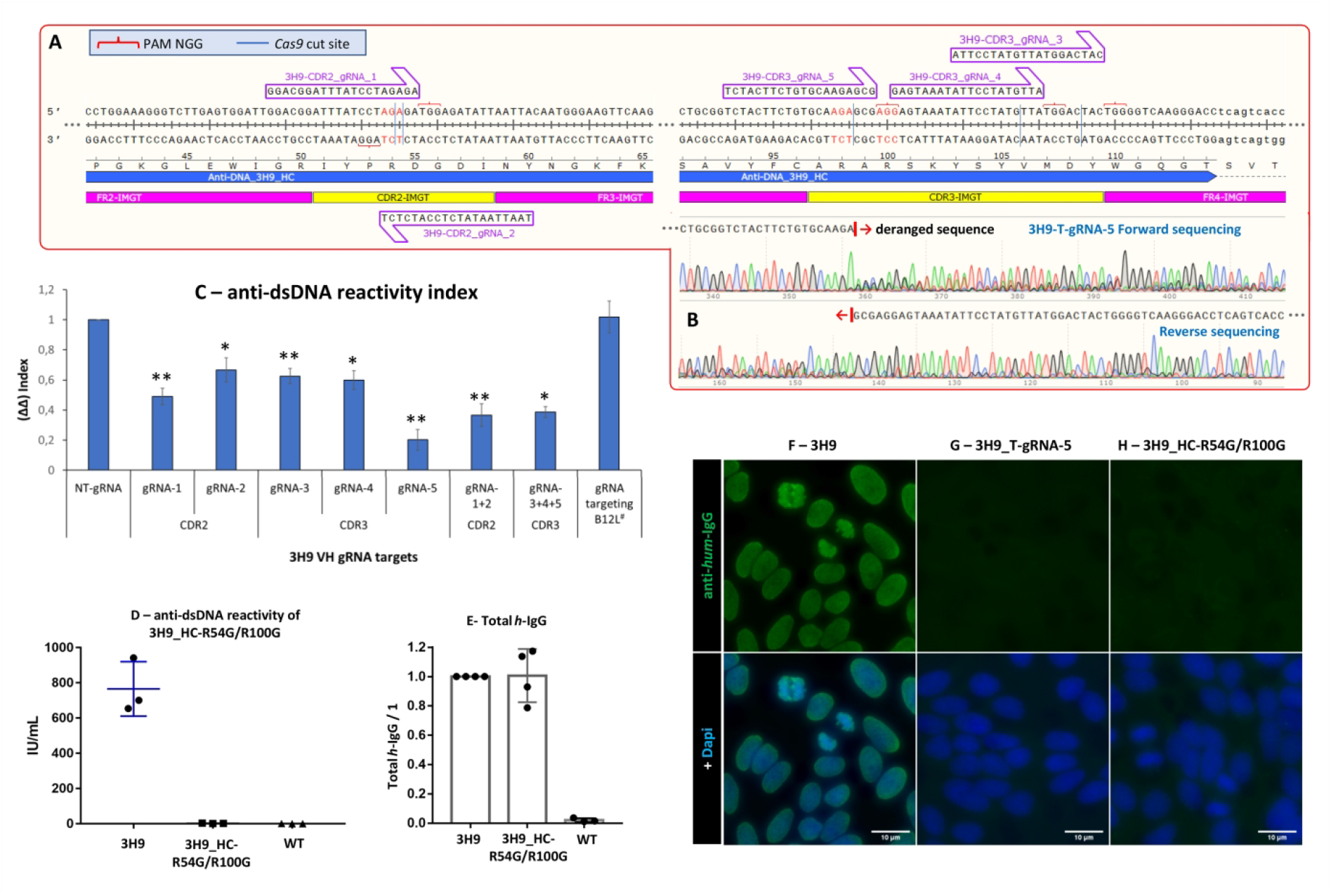
3H9 anti-dsDNA reactivity index in the recombinant antibodies produced after CRISPR editing. **(A)** Schematics showing 3H9-VH gRNAs targeting sites and CDR2/3 by IMGT. **(B)** After editing, DNA was extracted from the cells and the 3H9-VH region containing the gRNAs targeting sites was sequenced forward and reverse. This panel shows the sequence of 3H9-T-gRNA-5 targeting CDR3 with forward and reverse sequences deranged at the *Cas9* cut site. **(C)** ELISA levels of 3H9 anti-dsDNA International Units (IU/mL) divided by the total IgG levels (*h*-IgG in ng/mL), herein designated Δ index. In each batch of supernatant from cells subjected to CRISPR editing, the Δ index of T-gRNA was divided by the Δ index of NT-gRNA, generating the ΔΔ index. In the graph, the average ΔΔ index is shown for anti-dsDNA reactivity after editing with the 3H9 T-gRNAs, individually or grouped. Error bars = S.E.M. Statistics, values for each T-gRNA compared to NT-gRNA by the paired *t* test: *p<0.05; **p<0.01. **(D-E)** 3H9-VH antibody with the point mutations R54G and R100G (3H9_HC-R54G/G100G). **(D)** anti-dsDNA reactivity and **(E)** *h*-IgG levels measured in the supernatant of 3H9_HC-R54G/G100G producing cells. Error bars = S.D. **(F-H)** HEp-2-IFA using supernatant from **(F)** 3H9-HEK293T cells, **(G)** T-gRNA-5 knockout cells, and **(H)** 3H9_HC-R54G/G100G cells.

### 2.6. Genome editing also affects 3H9 anti-dsDNA reactivity

Since the knockout was not 100%, in addition to the levels of antibodies, we also evaluated if the CRISPR editing affects the 3H9 anti-dsDNA reactivity in the remaining antibodies produced by the cells. For the 3H9 antibody, both assays used to measure *h*-IgG (ng/mL) secretion levels and anti-dsDNA reactivity (IU/mL) were quantitative. Thus, by dividing the anti-dsDNA IU by the total *h*-IgG ng/mL (index Δ), we could estimate anti-dsDNA IU in each ng of 3H9. Among the four 3H9-NT-gRNA transfected cell supernatants, anti-dsDNA reactivity (index Δ) was ∼3.8 (SD ± 0.8), indicating that each ng of recombinant 3H9 provides near 4IU of anti-dsDNA reactivity. To analyze the effect of CRISPR editing on the anti-dsDNA reactivity of edited 3H9, we derived index ΔΔ by dividing index Δ of T-gRNA by index Δ of NT-gRNA in each batch of supernatant. This double index (ΔΔ) strategy allowed us to normalize anti-dsDNA reactivity per mass of *h*-IgG, and thus analyze if the CRISPR editing showed any effect over 3H9 dsDNA target binding in the remaining antibody produced by the transfected cells (Figure 4C).

The anti-dsDNA ΔΔ index was significantly lower in all 3H9-T-gRNAs edited antibodies compared to NT-gRNA (Figure 4C). For T-gRNAs 2, 3 and 4, the ΔΔ index was ∼0.6, indicating a ∼40% decrease in anti-dsDNA binding activity. For T-gRNA-1 alone, or the gRNAs targeting CDR2 grouped (1+2) or CDR3 grouped (3+4+5), the ΔΔ index was ∼0.5/0.4, indicating a ∼50-60% decrease in anti-dsDNA reactivity for these CRISPR editions. The lowest ΔΔ index was observed for 3H9-T-gRNA-5, ∼0.2, meaning a decrease of ∼80% in anti-dsDNA target binding activity for the remaining recombinant 3H9 antibody produced after the CRISPR editing with this specific T-gRNA (Figure 4C).

Previous reports have shown that Arginines (Arg/R) at CDR2 and CDR3 of the heavy chain of anti-dsDNA autoantibodies are important for antibody-antigen interaction [21-23], with higher relevance for the Arg in CDR3, not just the presence, but also its exact position at CDR3. In our 3H9-VH targeting strategy, T-gRNA-1 cut-site is at the Arg codon, position 54, in CDR2. At CDR3, the T-gRNA-5 cut-site is near the two Arg codons, positions 98 and 100, while T-gRNAs 3 and 4 cut-sites are over fifteen bases away from the Arg codon (schematics in Figure 4A). To further demonstrate the importance of Arg at 3H9-VH CDR2/3 for anti-dsDNA interaction, we generated a 3H9 antibody with two point-mutations, where Arg was replaced by Glycine (Gly/G) at positions 54 and 100 (3H9_HC-R54G/R100G). Without the Arg at the given positions, the recombinant antibody showed no anti-dsDNA reactivity (Figure 4D-H). Since anti-dsDNA reactivity in the remaining antibodies decreased ∼80% after T-gRNA-5 CRISPR/Cas9 editing, compared to ∼40% for T-gRNAs 3 and 4, we hypothesize that the *indels* introduced after Cas9 cut affected anti-dsDNA binding activity mainly by affecting Arg codons and, consequently, preventing translation of Arg at 3H9-CDR2/3.

## 3. Discussion / Conclusion

In this study, we generated two cell lines producing recombinant autoantibodies and demonstrated the reactivity to their respective targets: AChR and dsDNA. The variable regions of the heavy chain in each antibody were targeted by the CRISPR/Cas9 system, which knocked out the gene, resulting in ∼90% decrease in total IgG secreted. In addition, reactivity to the respective targets was also significantly affected in both antibodies, down to virtually undetectable levels for some T-gRNAs used in CRISPR knockout. Sequencing indicates the presence of *indels* at the Cas9 cut-site, which possibly led to codon jam, the likely cause of the knockout. However, in the remaining secreted antibodies after CRISPR editing, is likely that the *indels* introduced affected the translation of the heavy chain CDR by changing/adding/removing amino acids and further interfering with the antibody-antigen interaction.

In the design of the CRISPR/Cas9 system, several factors can affect gRNA efficiency in disturbing or knocking out the target gene: the presence of the protospacer adjacent motif (PAM), gRNA size and GC content, mismatches to avoid off-target effects, and other unpredictable facts [24]. For this study, we use the CHOPCHOP tool for gRNA design and choose five T-gRNAs for each antibody VH, two targeting CDR2 and three targeting CDR3. As expected, the 10 CRISPR/Cas9 T-gRNAs bound to, cut and knocked out their intended targets with various efficiencies. However, from ∼25% to ∼50% of the transfected cells still expressed some version of the VH-*h*IgG1-Fc_T2A_iRFP670 gene after CRISPR editing, as demonstrated by the reporter molecule fluorescence and the intracellular levels of IgG, although the secretion of measurable IgG decreased ∼90%. This difference could be explained by the fact that secreted antibodies will accumulate in the culture medium until the harvest, but not in the cell cytoplasm where they are readily secreted upon synthesis.

The efficiency of the different T-gRNAs varied, although their target sites are near to each other. In addition, conjugated transfection with different T-gRNAs showed no additive effect on the knockout. Since the targeted gene was a construct with a reported fluorescent molecule, we could purify cells with highly effective knockout, designated the “KO-Only” cells (Supp. Figure S4), which were sorted for the “absence” of the VH reporter iRFP670. These cells proved quite interesting, since secreted recombinant immunoglobulins were undetectable for these cells. Corresponding to the significant decrease in the secretion of IgG, reactivity to the respective targets by 3H9 and B12L also decreased, down to undetectable levels for some T-gRNAs, suggesting that the use of CRISPR/Cas9 systems to knockout autoantibodies could be a reasonable treatment strategy.

However, autoantibodies are polyclonal, although the repertoire of “disease-relevant” clones is limited. By using large databases of antibody sequences, such as cAb-Rep [25], AntiBodies Chemically Defined [26], and the international ImMunoGeneTics (IMGT) [27], among others, the repertoire of variable regions obtained from a given patient by next generation sequencing could be processed by artificial intelligence algorithms and machine learning to predict and choose a set of most relevant V(D)J sequences to be targeted by gRNAs, thus maximizing the efficiency in eliminating the relevant VH genes in plasma cell clones whose secreted autoantibodies present the most pathogenic potential.

Anti-dsDNA antibodies are highly specific, being found in 70–80% of SLE patients at some time during their illness [28]. Analysis of the VH sequence of over 500 anti-dsDNA antibodies from murine models and Systemic Lupus Erythematosus patients reveals that ∼70-80% present at least one arginine at the VH-CDR3 region [21, 22]. Furthermore, 35% of all anti-dsDNA antibodies present an arginine at position “96”, compared to only 5% in other conventional antibodies [22]. In our 3H9-VH antibody construct, position “96” corresponds to the arginine at position 100 (Figure 4A). Previous studies with the 3H9 antibody have demonstrated that mutation at the arginine in the VH-CDR3 region reduced drastically its affinity for dsDNA [23], indicating a possible hotspot for gRNA targeting and Cas9 cut-site in this study. Accordingly, anti-dsDNA reactivity in the 3H9 recombinant antibodies decreased ∼80% after T-gRNA-5 CRISPR/Cas9 editing near the two arginine codons at VH-CDR3 compared to ∼40% for T-gRNAs 3 and 4 with cut-sites over fifteen bases away from the arginine codons. We hypothesize that the *indels* introduced after the Cas9 cut could have affected arginine codons and, consequently, translation of arginine at the 3H9-CDR2/3 region, directly interfering with anti-dsDNA binding intensity.

Considerable progress in the treatment of autoimmune diseases has been made over the past 20 years with the advent of optimized immunosuppressive regimens and immunobiological drugs. Nevertheless, current therapies for autoimmune diseases still rely on non-specific modulation of the immune system. Therapies targeting the B cell lineage have little or no effect on long-lived plasma cells. Thus, side effects and suboptimal therapeutic results are still major problems in treating autoimmune diseases. In alignment with the concept of precision medicine, the therapy of autoimmune patients should address specific autoantigen systems and deranged immune-metabolic pathways at the individual level. This approach may be feasible with recent technologies, such as the Chimeric autoantibody receptor T cell (CAART), or the affinity matrix (AM), among others [11], including the CRISPR/Cas-based genome editing used in this study. With such approaches, the immune system could be modulated, suppressed, or enhanced, aiming to tolerate or attack specific autoantigens in the scenarios of autoimmunity and cancer, respectively. A successful example of antigen-specific modulation of the immune system is the active immunization with vaccines that has changed human health in the past century by preventing and eradicating dozens of severe infectious diseases.

In this proof-of-concept study, we tested if the CRISPR/Cas9 genome editing tool could be applied to knockout autoantibodies by targeting the epitope-specific sequences at the variable regions of the respective genes. As expected in our *in vitro* setup, CRISPR/Cas9 genome-editing was very effective in the knockout VH-IgG genes, considerably affecting the secretion of the recombinant autoantibodies *in vitro*. Additionally, the Cas9-mediated editing process significantly affected the ability of the remaining immunoglobulins to bind to their original targets, suggesting that the precise position of gRNA targeting and Cas9 cut-site is relevant also for eliminating reactivity to the original antigenic target. Adaptation of this method to *in vivo* modulation of pathogenic B/plasma-cell clones could pave the way to a novel antigen-specific targeted therapy in adherence to the concept of precision medicine.

## 4. Materials and methods

### 4.1. Plasmid cloning

#### 4.1.1. Plasmids for recombinant antibody production

PBMC were isolated from fresh human blood by density gradient (Ficoll-Paque PLUS 1.077 g/mL, Cytiva, USA). Total RNA was extracted with TRIzol (15596, Invitrogen, USA) and converted to cDNA with First Strand cDNA Synthesis Kit (E6300, NEB, USA). From the cDNA, the coding sequence of IgG1 constant part of the heavy chain (Reference sequence GenBank: MH975516.1) was amplified by PCR with Phusion Flash High-Fidelity PCR Master Mix (F548, Thermo Scientific, USA) using the primers shown in Supp. Table S1. The constant parts of the kappa light chain (Reference sequence GenBank: AF113887.1) and lambda light chain (Reference sequence GenBank: X57809.1) were amplified by the same method (Primers in Supp. Table S1).

PiggyBac inverted repeats for the transposon system were inserted in a plasmid with cytomegalovirus promoter (pCMV3) and hygromycin resistance gene (HygR) (Sino Biological, China). The following fragments were assembled into the linearized PiggyBac_pCMV plasmid: 1) *h*IgG1 + T2A + iRFP670 (fluorescent reporter gene Ex ∼643nm, Em ∼670nm); 2a) Constant L-Lambda + T2A + eGFP (fluorescent reporter gene Ex ∼488nm, Em ∼507nm); 2b) Constant L-kappa + T2A + eGFP. All fragments were assembled with NEBuilder HiFi DNA Assembly Cloning Kit (E5520S, NEB, USA), transformed into DH5-α Competent *E. coli* (C2987, NEB, USA), and the plasmids purified with Fast-n-Easy Mini-Prep Kit (DPK-104S, Cellco, Brazil). All gene sequences and assemblies were confirmed by Sanger sequencing.

The complete variable fragments, including the secretory signal peptides and the CDRs (∼500 bp each) for the anti-dsDNA 3H9 recombinant antibody (GenBank: M18234.1; M18237.1) and the B12L recombinant antibody [17] (obtained by personal communication with the study authors) were synthesized by FastBio Ltda, Ribeirao Preto, SP, Brazil and cloned into the respective vectors using the enzyme digestion sites *HindIII* and *AgeI* for plasmid 1 and *HindIII* and *XmaI* for plasmids 2a/b. Sequences of the variable fragments were confirmed by Sanger sequencing. The final plasmid maps are shown in Supp. Fig S1.

Point mutations were introduced through linearizing the plasmid by PCR with primers containing the intended mutations (Supp. Table S1) followed by reverting to circular plasmid with Gibson Assembly Master Mix (M5510A, NEB, USA).

#### 4.1.2. Plasmids for mAChR expression

Mouse muscle tissue *(Mus musculus)* was ground with a tissue homogenizer and RNA was extracted using TransZol Up Plus RNA Kit (ER501, TransGen Biotech, China). Reverse transcription to cDNA was performed with PrimeScrip RT Master Mix (RR036A, Takara Bio, Japan). The cDNA was used as a template for PCR amplification of the following gene coding sequences (primers shown in Supp. Table S1): AChR-α1 (NCBI: NM_007389.5); AChR-β1 (NCBI: NM_009601.4); AChR-δ (NCBI: NM_021600.3); AChR-γ (NCBI: NM_009604.3); AChR-ε (NCBI: NM_009603.1); Rapsyn (NCBI: NM_009023.3). The genes were cloned into the linearized pCMV3 vector (Sino Biological, China) by using ClonExpress UItra One Step Cloning Kit (C115, Vazyme, China) according to the manufacturer’s protocol. All gene sequences and assemblies were confirmed by Sanger sequencing.

Two final plasmids were constructed containing three genes each, interleaved by 2A sequences: 1) **pCMV_mAChR-α1_T2A_mAChR-β1_P2A_mAChR-δ-mTagBFP-Flag**, with the fluorescent reporter gene mTagBFP (Ex ∼399nm, Em ∼454nm) plus Flag tag fused to subunit δ; 2) **pCMV_mRapsyn_T2A_mAChR-γ_P2A_mAChR-ε_T2A_iRFP670-Myc**, with the fluorescent reporter gene iRFP670 fused to a Myc tag, segregated from the ε subunit sequence by a T2A.

#### 4.1.3. Plasmids for CRISPR editing

The designed gRNAs (Supp. Table S1) were cloned into a single guide RNA (sgRNA) expression vector. From the original plasmid pGL3-U6-sgRNA-EGFP (kindly donated by Xingxu Huang [29] #107721, Addgene, USA), the reporter gene was replaced with mTagBFP using the NEBuilder HiFi DNA Assembly Cloning Kit. Thus, the customized sgRNA expression vector was **pGL3-sp_U6-sgRNA_hPGK-mTagBFP**. For each gRNA, complementary forward and reverse oligos were synthesized and annealed. The vector was digested with Eco31I (BsaI) (ER0291, Thermo Scientific, USA) and the oligos were cloned into the BsaI site with T4 DNA Ligase (ENZ-101M, Cellco, Brazil). After transformation into DH5-α and Mini-Prep purification, the plasmids were sequenced to confirm the correct insertion of the sgRNA.

For the genome editing, we applied the spCas9 (Csn1) endonuclease from the *Streptococcus pyogenes* Type II CRISPR/Cas system. The original plasmid pST1374-NLS-flag-linker-Cas9 (kindly donated by Xingxu Huang [30] #44758, Addgene, USA), was modified to remove the flag linker. Thus, our final Cas9 expression vector was **pST1374_pCMV-Cas9-NLS**.

### 4.2. Cell culture and cell engineering

HEK293T cells were cultured in DMEM (31600-034, Gibco, USA), supplemented with 10% FBS (Nova Biotecnologia, Brazil), l-Glutamine 2mM and 1% Antibiotic–Antimycotic (A5955, Sigma, USA). Transfections were carried out with Lipofectamine 3000 (L3000-008, Invitrogen, USA) diluted in Opti-MEM I Reduced Serum Medium (22600-043, Gibco, USA), following the manufacturer’s protocol.

The antibody expression vectors for each recombinant antibody (VH + VL-L/k) were transfected together with a Transposase vector (Hera BioLabs, USA). Cells were selected with Hygromycin B (H3274, Sigma, USA) 100 μg/mL for ten days. Cells were then sorted for the presence of the reporter genes GFP and iRFP670 to generate a stable cell line secreting the recombinant antibodies.

For CRISPR editing, the antibody-producing cell lines were transfected simultaneously with the sgRNA expression vector and the spCas9 expression vector. Twenty-four hours after transfection, cells were sorted for the presence of the reporter gene mTagBFP. Five days later the cells were subjected to experiments for analyses of the effect of the CRISPR editing on the secreted antibodies.

For collecting of each batch of culture medium, antibody-producing cells (before or five days after the CRISPR/Cas9 editing) were resuspended with trypsin, washed once with PBS and seeded in 6-well plates at a density of 100.000 cells per well, in the presence of 2mL of FreeStyle 293 Expression Medium (12338018, Gibco, USA) supplemented with 1% Antibiotic–Antimycotic (Sigma, USA). After reaching 100% confluence, the culture medium was collected, filtered in 0.22μm syringe filters, and stored at -20°C for analysis of the secreted antibodies. The remaining cells attached to the plate were collected for DNA and protein analysis.

### 4.3. Cytometry analysis and cell sorting

For analysis of the reporter genes, cells were resuspended with trypsin, washed once with PBS, and analyzed in a CytoFLEX cytometer (Beckman Coulter, USA). One hundred thousand valid events gated from FSC-SSC were collected for each sample analyzed. For cell sorting, cells were resuspended with trypsin, washed once with PBS, and resuspended in 1ml of FACS buffer (PBS pH7.2, BSA 1%, EDTA 2mM, HEPES 20mM, Sodium Azide 0.05%). The sorting was carried out in a BD FACSAria II Cell Sorter (Becton Dickinson, USA). Fifty thousand cells expressing the reporter gene were sorted for each sample.

### 4.4. DNA extraction and sequencing after the CRISPR editing

Total DNA was extracted from the cells with FlexiGene DNA Kit (51206, Qiagen, Germany) following the manufacturer’s protocol. The obtained DNA was quantified using a NanoDrop ND-1000 Spectrophotometer (Thermo Fischer, USA) and stored at -20°C. The variable region of the heavy chains of 3H9 and B12L, containing the CDR2/3 gRNA targeting sites, were amplified by PCR with Phusion Flash Master Mix (Thermo Scientific, USA) and specific primers (Supp. Table S1). The fragment (∼560bp) quality was evaluated in 2% agarose gel electrophoresis, before sequencing by the Sanger method using the forward and reverse primers.

### 4.5. Protein extraction and Western Blot

For total protein extraction, cells were ressuspended with trypsin and washed once with 1mL PBS. From each well of a 6-well plate, the pellet was lysed with 100μL of RIPA buffer (Tris 50mM, NaCl 15mM, 1% IgepalCA-630, Sodium deoxycholate 0.5%, EDTA 1mM, SDS 0.1%), plus 10μL of Halt Protease inhibitor Cocktail (1862209, Thermo Scientific, USA). After disruption by sonication, the material was centrifuged at 10.000rpm and the supernatant was stored at -20°C.

Both protein cell extract and the collected culture medium were analyzed independently by western blot. Samples were denatured at 90°C for 10min in the presence of 4x LDS Sample Buffer (NP0007, Invitrogen, USA) and 10x Sample Reducing Agent (B0009, Novex, USA). 10μL of the sample was loaded onto each well of the gel (NuPAGE 4-12% Bis-Tris Gel, Invitrogen, USA), run with XCell SureLock Mini-Cell Electrophoresis System and transferred to a 0.45 Micron nitrocellulose membrane (88025, Thermo Scientific, USA) with XCell SureLock Blot Module (Thermo Fisher Scientific, USA). After transfer, the membrane was incubated with agitation for 1h with blocking buffer (Tween 0.1%, skim milk 5% in PBS) at room temperature.

For immunolabeling, the antibodies were diluted in blocking buffer and incubated overnight at +4°C with agitation. The membrane containing the collected culture medium was incubated with goat anti-human IgG conjugated to IRDye 800CW (926-32232, Li-Cor, USA) diluted at 1:3000 and the near-infrared fluorescence signal was analyzed by an Odyssey CLx scanner (Li-Cor, USA). The membrane containing the total cell extract was incubated with HRP-conjugated anti-GAPDH mouse monoclonal antibody (HRP-60004, ProteinTech, USA). The HRP signal was revealed by ChemiFast Chemiluminescent Substrate (CH-FAST/20, Syngene, UK) and visualized by an imaging system (UVITEC 4.7, Cambridge, UK).

### 4.6. ELISA

Total human IgG in the collected culture medium was measured by the quantitative SimpleStep ELISA Kit (ab195215, ABCam, UK) following the manufacturer’s protocol. Samples were applied diluted 1/40 as recommended by the manufacturer, since this kit has high sensitivity for low concentration of human IgG, down to ng/mL.

For analysis of intracellular human IgG, cells were resuspended with trypsin and washed once with 1mL PBS. Each well of a 6-well plate was incubated for 30min at room temperature with 200μL of Cell Extraction Buffer PTR (ab195215, ABCam, UK) containing phosphatase inhibitors and the protease inhibitor aprotinin. After disruption by sonication, the material was centrifuged at 10.000rpm to remove debris, and the amount of protein was quantified by Bradford assay [31] using a colorimetric Protein Assay Dye (500-0006, Bio-Rad, USA) and Bovine Gama Globulin in known concentration for the standard curve. Samples diluted 1/10 were applied to each well of the ELISA plate (ab195215, ABCam, UK) and the protein concentration in each sample was normalized to the NT-gRNA protein concentration in the same run.

Anti-dsDNA reactivity for the recombinant IgG antibodies in the supernatant was measured using a quantitative anti-dsDNA-NcX ELISA (EA-1572-9601-G, Euroimmun, Germany) following the manufacturer’s protocol. Samples were applied diluted 1/2 and the obtained optic density (O.D.) interpolated in the calibration curve to find anti-dsDNA reactivity expressed in International Units (IU/mL).

### 4.7. Indirect Immunofluorescence

For B12L anti-AChR analysis by indirect immunofluorescence, the HEK293T cells expressing mouse AChR (HEK293T-mAChR) and wild-type HEK293T cells were cultured to ∼80% confluence in 13mm round coverslips. After washing with PBS, cells were fixed with 4% paraformaldehyde for 15 minutes and extensively washed in PBS. Collected supernatant from B12L-expressing cells was added diluted 1/10 in PBS + BSA 1%, and incubated overnight at +4°C in a moist chamber. To preserve cell membrane stability, and therefore the AChR integrity, the detergent-based permeabilization step was omitted. After 3 washes with PBS, cells were incubated for 1h with anti-human IgG goat antibody conjugated to FITC or Alexa Fluor 488 (H10301; A-11013, Invitrogen, USA) diluted 1/500 in PBS. After additional 3 washes, coverslips were mounted with VECTASHIELD Mounting Medium with DAPI (VECTOR Labs, USA), sealed and analyzed with a fluorescence microscopy system (Axio Imager.M2, Carl Zeiss, Germany).

3H9 reactivity to dsDNA was tested by indirect immunofluorescence using commercial HEp-2-IFA slides as substrate (FA-1520-2010, Euroimmun, Germany). The procedure followed the manufacturer’s protocol, except for the dilution of the collected supernatants, which started at 1/10 followed by sequential double dilutions, 1/20, 1/40 … and so on. Slides were also mounted with VECTASHIELD Mounting Medium with DAPI, sealed and analyzed in fluorescence microscopy.

### 4.8. Determination of anti-AChR reactivity of B12L antibodies by flow cytometry

For evaluation of B12L anti-AChR reactivity by flow cytometry, the immunolabeling reaction was performed using alive (non-fixed) HEK293T cells expressing mouse AChR (HEK293T-mAChR) in suspension. Briefly, HEK293T-mAChR cells were resuspended with trypsin, followed by incubation with culture supernatant collected from B12L-expressing cells diluted 1/4 in the incubation medium (DMEM supplemented with BSA 1%, EDTA 5mM, HEPES 20mM and sodium azide 0.05%) for 1h at 37°C under gentle agitation. After centrifugation, cells were washed once by resuspending the pellet with 500μL of FACS buffer, followed by fixation with 4% paraformaldehyde for 15 min. After another washing, cells were incubated for 1h with anti-human IgG goat polyclonal antibodies conjugated to FITC or Alexa Fluor 488 (H10301; A-11013, Invitrogen, USA) diluted 1/100 in FACS buffer. After two final washing steps, the pellet was resuspended in 500μL of FACS buffer and analyzed in a CytoFLEX cytometer (Figure 2A-C).

### 4.9. Data analysis

All cytometry data were analyzed with CytExpert v2.3 software, with 30k to 50k events shown in the graphs, unless indicated otherwise. Western blot experiments with the culture supernatant were performed in quadruplicate and experiments with the total cell extract in duplicate. The intensity of the IgG band was quantified using the Surface Plot tool in ImageJ software, normalized by the average of GAPDH as a reference. Plasmid construction was designed and DNA sequencing results were analyzed with SnapGene v3.2.1 software.

The data are presented as mean plus error bars indicating Standard Deviation (S.D.) or Standard Error of the Mean (S.E.M.) as described in figure legends. For some analyses, such as total IgG levels and antibody reactivity, the IgG concentration or antibody reactivity of the supernatant of T-gRNAs transfected cells was normalized to the values obtained from the supernatant of NT-gRNA transfected cells in each batch of culture medium collected, thereby allowing to estimate the amount of variation in T-gRNA transfected cells. For statistical analysis, original values (without normalization to 1) for each T-gRNA were compared to NT-gRNA, using the paired *t*-test. p ≤ 0.05 was considered statistically significant. All analyses were done with GraphPad Prism v7.0 software.

## 5. Author statements

### Research Funding

This work was supported by Sao Paulo Government agency FAPESP (Sao Paulo State Research Foundation) grant numbers #2017/20745-1 and #2021/04588-9, granted to G.D.K. and L.D.S. Additionally, L.E.C.A. is supported by the Brazilian research agency National Council for Research (CNPq), grant #PQ-1D 310334/2019-5.

The funding organizations played no role in the study design; in the collection, analysis and interpretation of data; in the writing of the report; or in the decision to submit the report for publication.

### Authors contribution

G.D.K. conceived the study, designed and performed the experiments and wrote the manuscript; L.D.S. and K.G.S. performed experiments and interpret data; L.E.C.A. conceived the study, designed the experiments, and critically reviewed and edited the manuscript.

All authors have accepted responsibility for the entire content of this manuscript and approved its submission.

### Ethical approval

The study was approved by the local Ethics Committee at the Federal University of Sao Paulo (Plataforma Brasil CAAE: 08170919.1.0000.5505).

## Acknowledgments

We acknowledge Tomohiro Makino, Ph.D. and co-authors for kindly sharing, by personal communication, the sequence for the variable region of the B12L recombinant antibody.

We acknowledge the team in the Flow Cytometry Facility, especially Dr. Daniela Teixeira and Prof Alexandre Basso, from “Departamento de Microbiologia, Imunologia, e Parasitologia, Escola Paulista de Medicina” at the Federal University of Sao Paulo - UNIFESP, for the assistance with cell sorting.

## Data Availability

The datasets generated during and/or analyzed during the current study are available from the corresponding author upon reasonable request.

