## Supplementary material for "“UNTARGETING” AUTOANTIBODIES USING GENOME EDITING, A PROOF-OF-CONCEPT STUDY": Supp. Figure

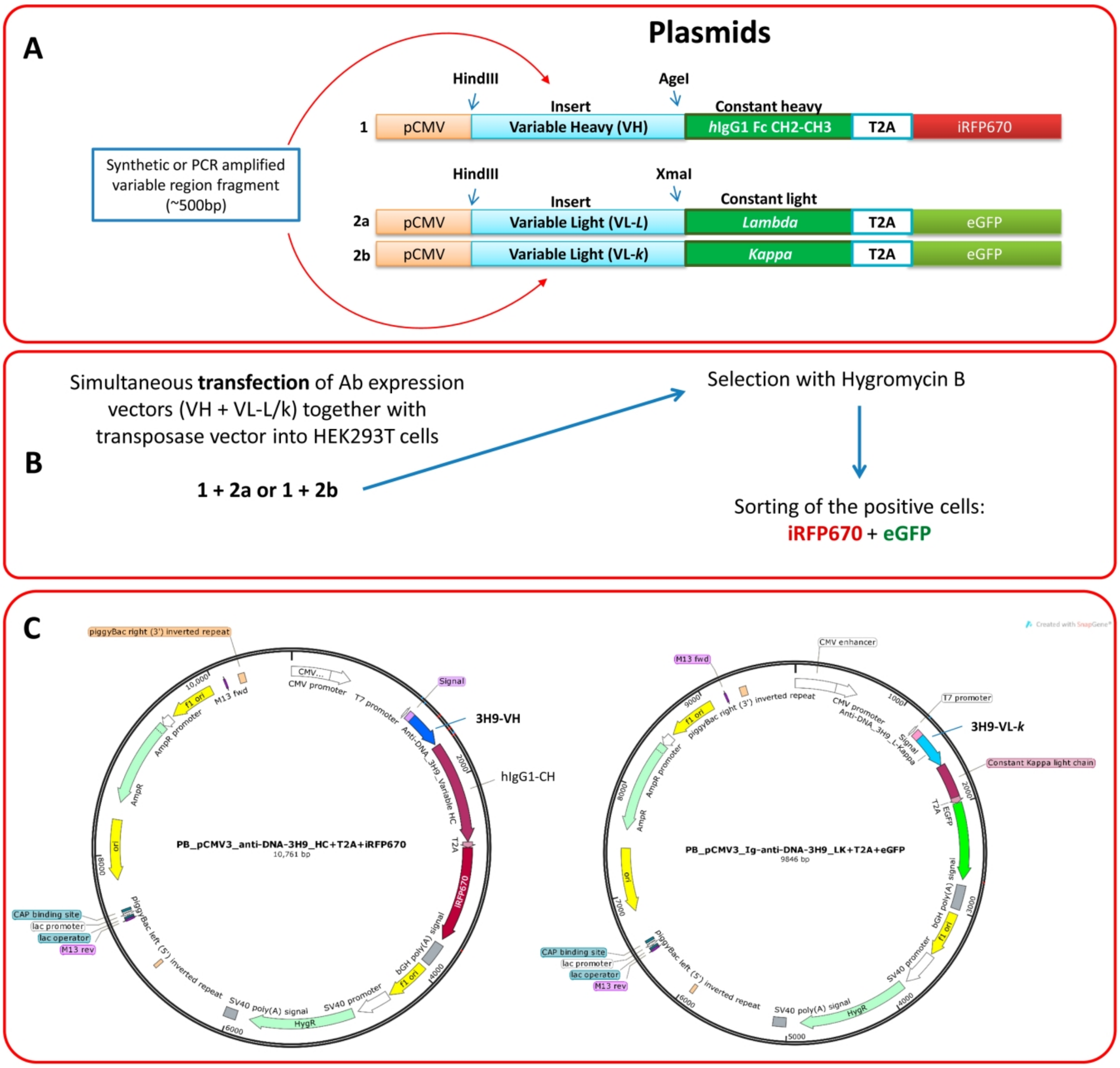

**Supplementary Figure S1. Strategy for construction of plasmids and antibody-expressing cell-lines. (A)** Plasmid 1 contains the human immunoglobulin constant region of the Heavy Chain, plus a reporter fluorescent gene (iRFP670) interleaved by T2A. Plasmid 2 contains the constant Light Chain plus eGFP, interleaved by T2A. Plasmid 2a is Light Lambda and 2b is Light kappa. The variable region of the desired antibody could be obtained by synthesis or by PCR amplification. After enzymatic digestion, the fragment was inserted into the vectors. **(B)** Both VH and VL plasmids for recombinant IgG synthesis were transfected into the cells, together with a Transposase vector. After Hygromycin selection and sorting of the cells expressing the reporter genes, a cell line was established secreting the desired recombinant human antibody. **(C)** Plasmid maps for 3H9 recombinant antibody as an example, containing pCMV and the PiggyBac inverted repeats.

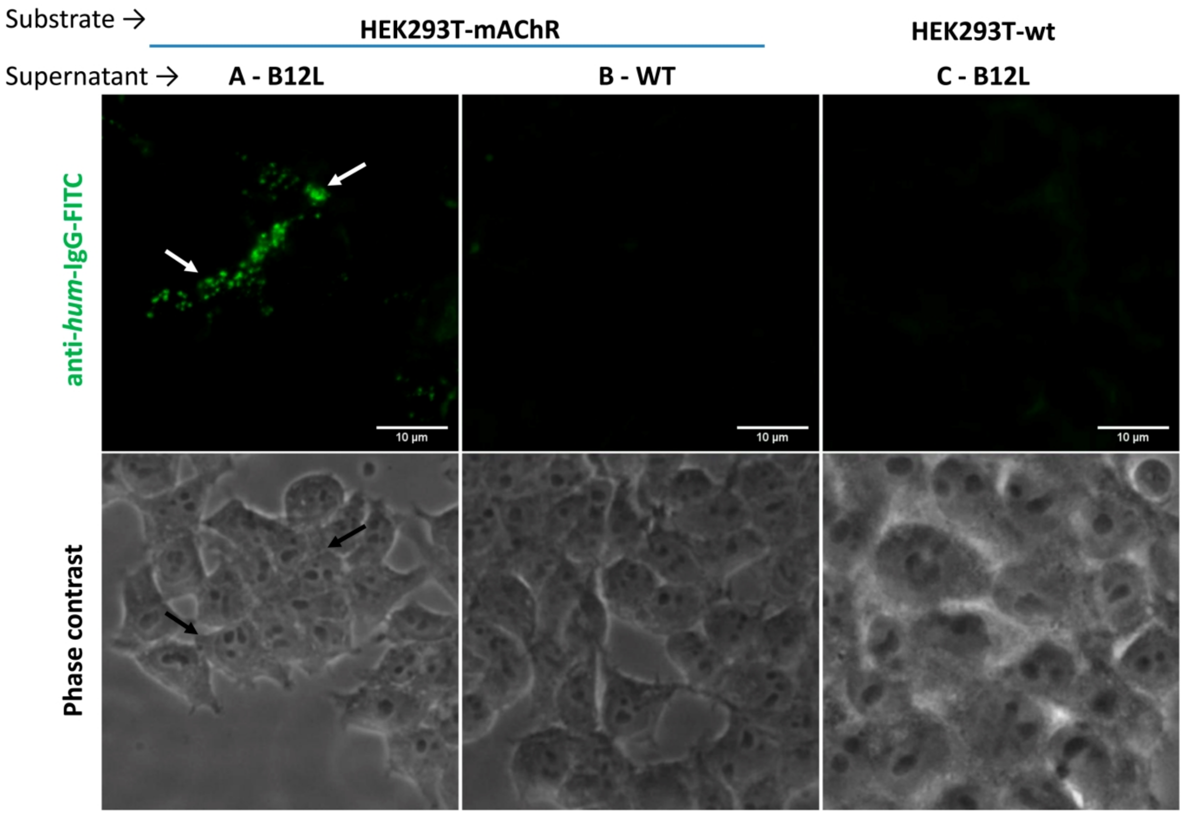

**Supplementary Figure S2. Cells expressing the mouse AChR-α1/β1/δ/γ/ε-subunits as substrate for indirect immunofluorescence (IIF) assay. (A-B)** HEK293T-mAChR or wild-type cells were used as substrate in an IIF with supernatant containing recombinant antibody as primary probe collected from B12L antibody-producing cells **(A)** or control "non-transfected" wild-type (WT) cells **(B)**. **(C)** When wild-type HEK293T was used as substrate, supernatant from B12L antibody-producing cells labels nothing. Arrows in **(A)** indicate AChR clusters, labeled by the recombinant B12L (green). The bottom row shows the field phase contrast images for visualization of the cells contour.

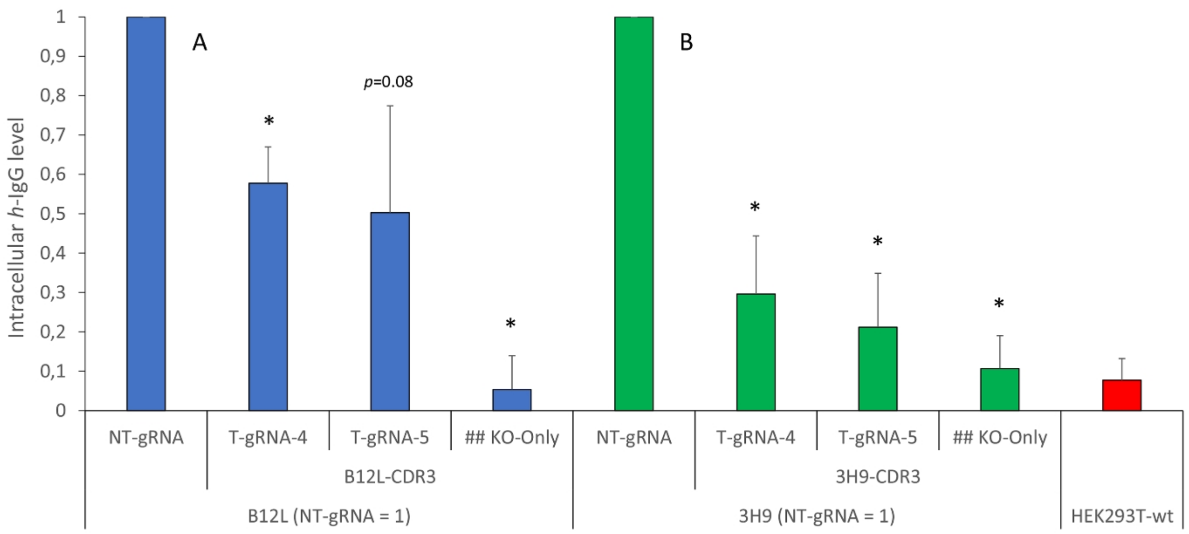

**Supplementary Figure S3. Intracellular IgG levels after CRISPR/Cas9 editing.** Total cell extract was collected after CRISPR editing and total *h*-IgG was analyzed by ELISA. **(A)** B12L T-gRNA knockout and **(B)** 3H9 T-gRNA knockout. For both analysis, concentration in NT-gRNA was considered as 1 in each batch of culture medium collected (n=4). Error bars = S.D. Statistics, values for each T-gRNA compared to NT-gRNA by Paired *t* test: *p<0.05.

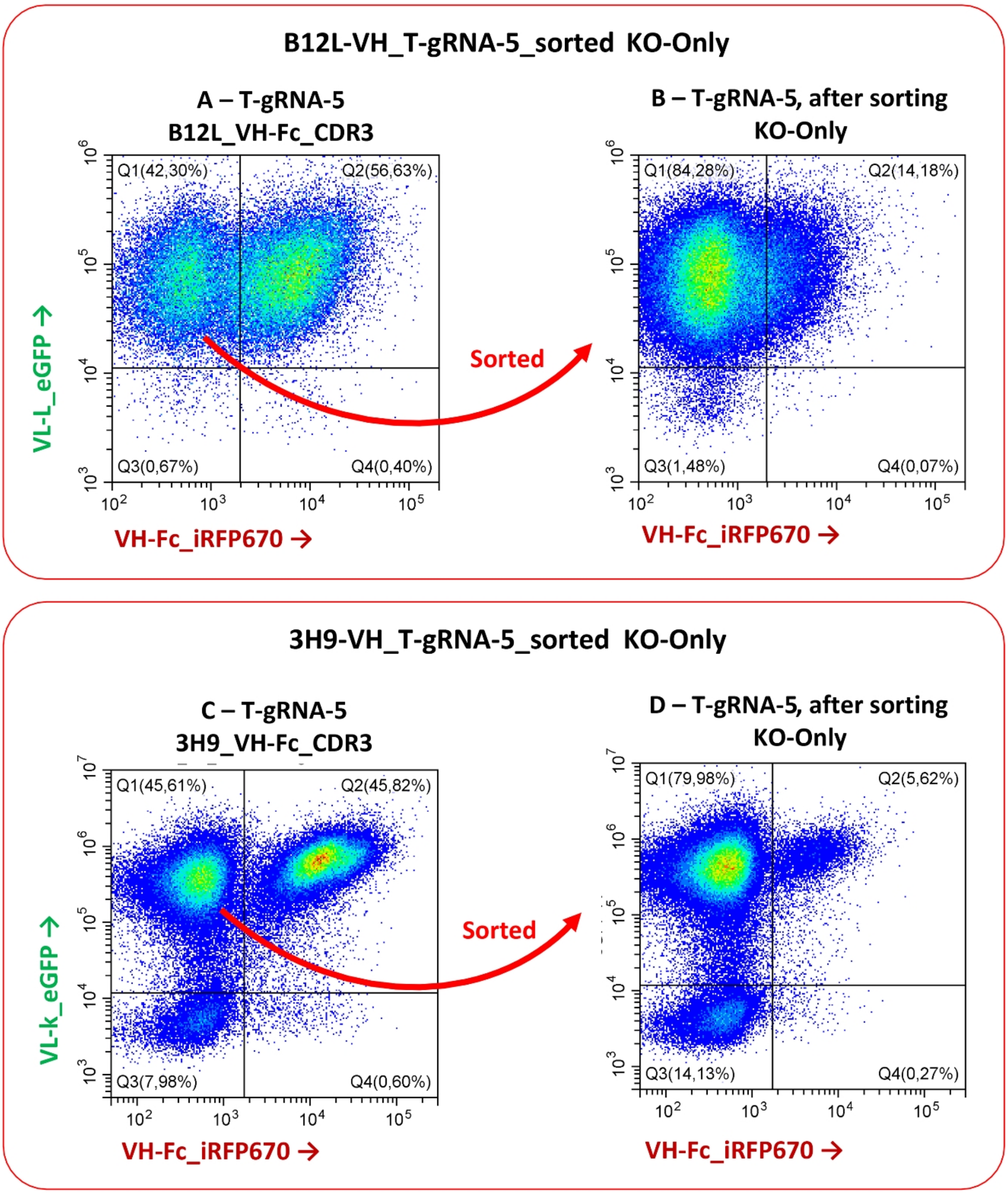

**Supplementary Figure S4. Sorting of knockout-only cells.** Five days after CRISPR/Cas9 editing, the cells that "lost" expression of the VH-Fc reporter gene (iRFP670) were sorted, resulting in a population of cells where the VH-Fc CRISPR knockout was "highly effective", we name this population "KO-Only" cells. **(A-B)** Example of T-gRNA-5 targeting B12L-VH-Fc, sorting to generate B12L-KO-Only cells. **(C-D)** Example of T-gRNA-5 targeting 3H9-VH-Fc, sorting to generate 3H9-KO-Only cells.

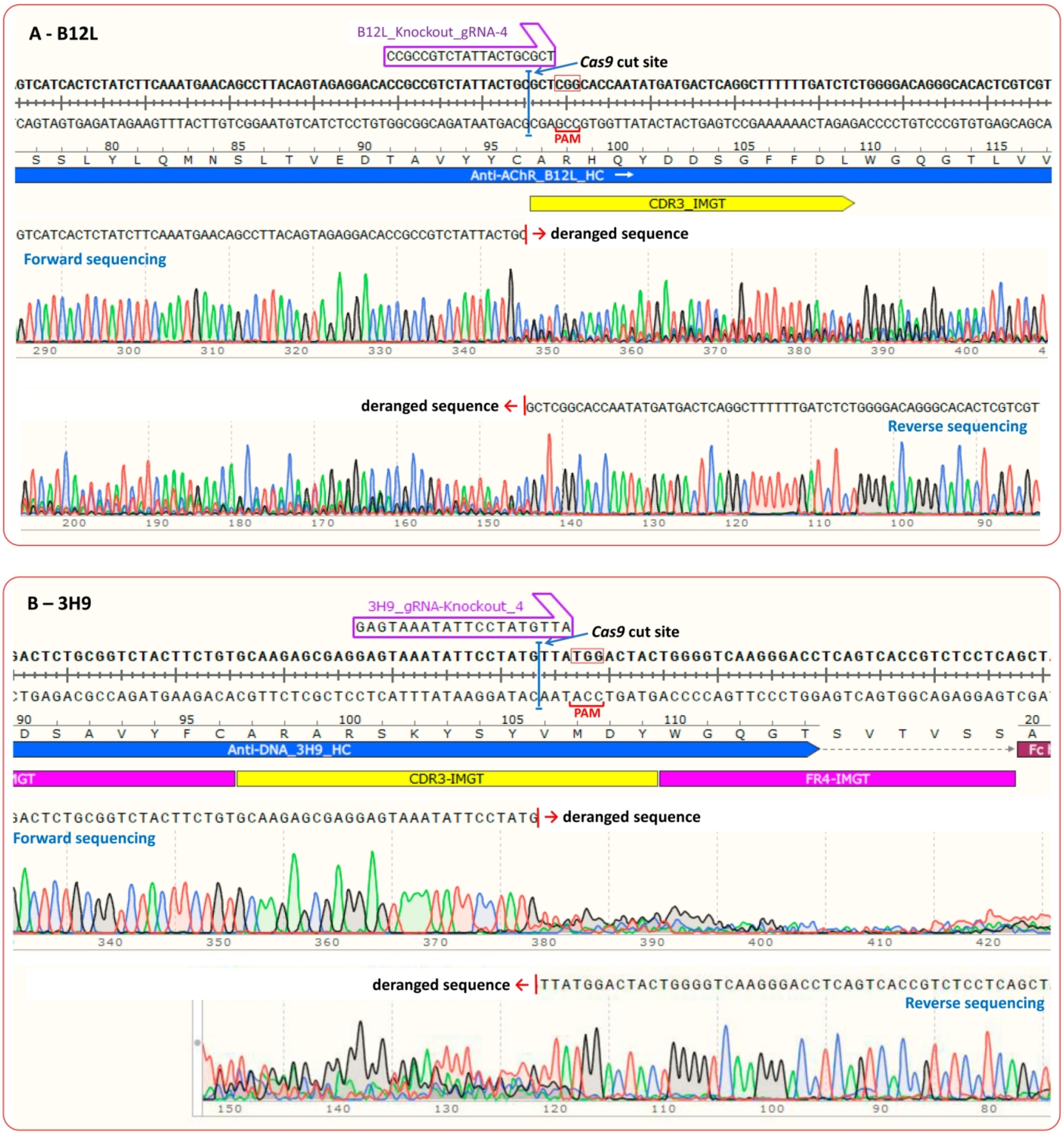

**Supplementary Figure S5. VH sequencing after CRISPR editing.** After CRISPR-Cas9 editing, DNA was extracted from the B12L-expressing cells and from the 3H9-expressing cells. The Heavy Chain variable regions containing the T-gRNA sites were sequenced forward and reverse. **(A)** In this figure, T-gRNA-4 targeting B12L-VH-Fc CDR3 is shown as example. **(B)** In this figure, T-gRNA-4 targeting 3H9-VH-Fc CDR3 is shown as example. Forward and reverse sequencing were deranged at *Cas9* cut site for both antibodies, indicating the presence of *indels*.

**Supplementary Table S1. Primers and gRNA sequences** (shown in 5’-3’).

| For amplification of coding sequences | | Forward | Reverse |
|---|---|---|---|
| human PBMC cDNA | IgG1 constant Heavy chain | GCCTCCACCAAGGGCCCATCGGTCTTC | CTTGCCGGGGGACAGGCTCAGGCTCTTC |
| | Constant Light-kappa | CGAACTGTGGCTGCACCATC | GCACTCTCCCCTGTTGAAGC |
| | Constant Light-Lambda | GGTCAGCCCAAGGCTGCCCC | TGAACATTCTGTAGGGGCCAC |
| Mouse muscle cDNA | AChR-α1 | ATGGAGCTCTCGACTGTTCTCC | TCCTTGTTGATGTAACTCAATG |
| | AChR-β1 | ATGGCCTTAGGGGCGCTGCTTC | AGGGAAAGGTTCAGGAGGGGGC |
| | AChR-δ | ATGGCAGGGCCTGTGCTCACAC | GATGAAGCGCTTGTCCTGTTCG |
| | AChR-γ | ATGCAAGGGGGCCAGAGACCTC | GTCTGGCAAAGGCAGGTAGGGG |
| | AChR-ε | ATGGCAGGGGCTCTGCTTGGTG | TGGTTGGATGCACGGTGGGTAAG |
| | Rapsyn | ATGGGGCAGGACCAGACAAAGC | CACAAAGCCCGGCTTCATGGAG |
| | | gRNA targeting VH-Fc (for *Cas9* knockout) (NGG_PAM) | |
| | | **B12L** | **3H9** |
| T-gRNA-1 | | TACGGATACGCCGTATCTAG TGG | GGACGGATTTATCCTAGAGA TGG |
| T-gRNA-2 | | AGATACGGCGTATCCGTAAA AGG | TAATTAATATCTCCATCTCT AGG |
| T-gRNA-3 | | GTGCCGAGCGCAGTAATAGA CGG | ATTCCTATGTTATGGACTAC TGG |
| T-gRNA-4 | | CCGCCGTCTATTACTGCGCT CGG | GAGTAAATATTCCTATGTTA TGG |
| T-gRNA-5 | | CGGCACCAATATGATGACTC AGG | TCTACTTCTGTGCAAGAGCG AGG |
| NT-gRNA | | TGAGACCGAGAGAGGGTCTCA | |
| For VH amplification and sequencing after the CRISPR editing | | **Forward** | **Reverse** |
| | | TAATACGACTCACTATAGGG | GTTCGGGGAAGTAGTCCTTGAC |
| **Point-mutations** | 3H9_VH-R54G | TCCT**G**GAGATGGAGATATTAATTACAATGGG | AATATCTCCATCTC**C**AGGATAAATCCGTCC |
| | 3H9_VH-R100G | GAGCG**G**GGAGTAAATATTCCTATGTTATGGAC | GGAATATTTACTCC**C**CGCTCTTGCACAGAAGT |
